# Transmission of synthetic seed bacterial communities to radish seedlings: impact on microbiota assembly and plant phenotype

**DOI:** 10.1101/2023.02.14.527860

**Authors:** Marie Simonin, Anne Préveaux, Coralie Marais, Tiffany Garin, Gontran Arnault, Alain Sarniguet, Matthieu Barret

## Abstract

Seed-borne microorganisms can be pioneer taxa during germination and seedling emergence. Still, the identity and phenotypic effects of these taxa that constitute a primary inoculum of plant microbiota is mostly unknown. Here, we studied the transmission of bacteria from radish seeds to seedlings using the inoculation of individual seed-borne strains and synthetic communities (SynComs) under *in vitro* conditions. The SynComs were composed of highly abundant and prevalent, sub-dominant or rare bacterial seed taxa. We monitored the transmission of each strain alone or in communities using *gyrB* gene amplicon sequencing and assessed their impacts on germination and seedling phenotype.

All strains and SynComs successfully colonized seedlings and we were able to reconstruct a richness gradient (6, 8 and 12 strains) on both seeds and seedlings. *Stenotrophomonas rhizophila* became dominant on seedlings of the three SynComs but most strains had variable transmission success (i.e increasing, stable or decreasing during seed to seedling transition) that also depended on the SynCom richness.

Most individual strains had no effect on seedling phenotypes, at the exception of *Pseudomonas viridiflava* and *Paenibacillus sp. that* had detrimental effects on germination and seedling development. Abnormal seedling morphologies were also observed with SynComs but their proportions decreased at the highest richness level. Interestingly, some bacterial strains previously identified as core taxa of radish seeds (*Pseudomonas viridiflava, Erwinia persicina)* were associated with detrimental effects on seedling phenotypes either in isolation or in SynComs. These results confirm that the plant core microbiome includes pathogenic and not only commensal or mutualistic taxa.

Altogether, these results show that SynCom inoculation can effectively manipulate seed and seedling microbiota diversity and thus represents a promising tool to better understand the early stages of plant microbiota assembly. This study also highlights strong differences between native seed-borne taxa in the colonization and survival on plant habitats.

## INTRODUCTION

The impact of seed-borne pathogens on plant fitness has been extensively studied but the influence of all the other commensal or mutualistic microorganisms living on/in seeds is mostly unknown. Seed microbiota harbor diverse microbial communities that constitute a primary inoculum for plant microbiota assembly that could originate from maternal transmission or environmental sources during seed maturation (Chesneau *et al*. 2020, 2022). Several individual seed endophytic bacteria, such as methylotrophic taxa, have been reported to promote stress tolerance and germination (Kumar *et al*. 2019, Raja *et al*. 2019). However, the research is still scarce on the influence of complex seed microbial communities on germination and seedling phenotypes or on the role of microbial interactions in the observed phenotypes (Lamichhane *et al*. 2018). Moreover, many knowledge gaps remain regarding the fraction of seed-borne taxa that can be transmitted to seedlings and the influence of the initial seed microbiota composition on their transmission success.

Few studies investigated the transmission of seed microbiota to seedlings and they reported a high variability in the contribution of seed taxa to plant microbiota depending on the soil tested (Rochefort *et al*. 2021; Walsh *et al*. 2021). These studies had the advantage to work directly on native seed communities but this comes with the limitation that the seed microbiota had to be characterized on pools of seeds due to the insufficient microbial biomass and DNA present on individual seeds. As a consequence, it is not possible to truly determine which microbial taxa were transmitted from one seed to one seedling. Few studies attempted to characterize the microbiota of individual seeds using culture-dependent (Mundt and Hinkle 1976; Newcombe *et al*. 2018) and culture-independent techniques (Bintarti *et al*. 2021; Chesneau *et al*. 2022). These studies demonstrated a very high natural variability in microbiota composition between individual seeds and a low bacterial richness. In particular, our work on radish seeds showed a high variability in richness (1 to 46 bacterial taxa, Figure S1) and composition between individual seeds (Chesneau *et al*. 2022). An important finding was that seeds were generally dominated by one bacterial taxon (> 75% relative abundance) but its identity was highly variable between parental plants and even between seeds originating from the same plant.

Hence, studying the transmission of natural microbiota from seed to seedling and assessing their impacts on seedling phenotype is extremely challenging. Controlled conditions with the inoculation of known microbial assemblages are required to characterize the seed to seedling transmission and establish causal links between microbiota and seedling phenotypes.

The contribution of the plant microbiota to host nutrition or resistance to pathogens has been recently investigated through inoculation of synthetic microbial communities (SynComs) in the rhizosphere and the phyllosphere (e.g. Kwak *et al*. 2018; Carlström *et al*. 2019; Finkel *et al*. 2020). But so far, assembly and role of the plant microbiota in the early stage of the plant’s life cycle has been neglected. One correlative study between seed microbiota structure and seed germination of different rapeseed genotypes offers promising results about the key role of microbiota in seed vigor (Rochefort *et al*. 2019). One SynCom study by Figueiredo dos Santos *et al*. (2021) showed that seed disinfection reduced maize germination rates that could be recovered after inoculation of a SynCom on seeds. Additionally, Matsumoto et al. (2021) demonstrated the role of seed endophytes in disease resistance to the seed-borne pathogen *Burkholderia plantarii*. These reports encourage further investigations on the influence of seed microbiota on seedling microbiota assembly to uncover the role of this primary inoculum of plants.

In this context, we set up seed inoculation experiments under *in vitro* conditions to address the following objectives:

- Monitor the transmission of individual seed-borne bacteria and synthetic bacterial communities from seed to seedlings (here on radish plants) using *gyrB* amplicon sequencing
- Test the hypothesis that individual seed-borne bacteria and/or synthetic bacterial communities have significant effects on seedling phenotype

Twelve bacterial strains representative of radish seed microbiota were selected based on their abundance – prevalence in seeds without any *a priori* on their potential functions or effects on plants. The strains were studied either individually or in communities (mix of 6, 8, or 12 strains). In this study, we considered the entire seed and seedling microbiota, including both endophytes and bacteria living on the plant surfaces.

## MATERIAL AND METHODS

### Seed material and bacterial strains

The radish seeds (*Raphanus sativus* var. Flamboyant5) used for bacterial strain isolation and the inoculation experiments were obtained from a field trial conducted in 2013 and 2014 at the experimental station of the National Federation of Seed Multipliers (FNAMS, 47°28’012.42’’N - 0°23’44.30’’W, Brain-sur-l’Authion, France).

A bacterial culture collection was obtained from a seed sample composed of approximately 1000 mature seeds (Torres-Cortés et al. 2019). Seed samples were soaked in 25 ml of phosphate-buffered saline (PBS, Sigma-Aldrich) with Tween® 20 (0.05 % v/v, Sigma-Aldrich) at 6°C under agitation (150 rpm) for 2h30. This seed soaking technique has been developed by the International Seed Testing Association for the detection of bacterial pathogens of Brassicaceae. This technique permits isolation and characterization of both the endophytic and epiphytic fractions of the radish seed microbiota (Chesneau et al. 2022). One hundred microliters of the suspension were plated on tryptic soy agar 1/10 strength (10% TSA) supplemented with cycloheximide (50 µg.ml-1, Sigma-Aldrich, Saint-Louis, Missouri, USA). Plates were incubated at 20°C during a minimum 5 days. Isolated colonies were picked and grown on 10% TSA plates for 24-48 hours to obtain pure cultures. The taxonomy of the isolated strains was determined using Sanger sequencing of the *gyrB* gene. A total of 528 strains were obtained that represented the 4 main phyla (Proteobacteria, Bacteroidetes, Firmicutes, Actinobacteria) found in radish seeds, for a total of 11 assigned bacterial families and 17 genera.

To select strains representative of the radish seed bacterial community, we used the Seed Microbiota Database (Simonin *et al*. 2022) originating from a meta-analysis that gathers re-processed amplicon sequencing datasets from 50 plant species. All the radish seed datasets (*R. sativus* Flamboyant5) obtained with the *gyrB* marker gene (7 independent studies, n=295 seed samples) were extracted from the database. After filtering the ASVs with a low read number (<100 reads), the prevalence and relative abundance of ASVs (n=139) were calculated across all samples. The *gyrB* sequences of the ASVs were compared to the strains of our culture collection (Supplementary Data).

### Seed inoculation experiment

Subsamples of seeds (1g ∼100 seeds) were surface-sterilized using the following protocol: 1 min sonication (40 Hertz), soaking for 1 min in 96° ethanol, 5 min in 2.6% sodium hypochlorite, 30 sec in 96° ethanol and rinsed 3 times with sterile water. A subsample of 30 seeds was used to verify the efficacy of the surface-sterilization by soaking the seeds under agitation (150 rpm) for 2h30, plating on 10% TSA and incubating a minimum of 2 days at 20°C. The seeds were dried on sterile paper before the inoculation. Of note, the surface-sterilization cannot eliminate the seed endophytes without impairing seed physiology and germination potential. Endophyte diversity was therefore estimated in non-inoculated control seeds and seedlings.

Twelve bacterial strains of our culture collection conserved at the CIRM-CFBP were selected to build a simplified seed radish bacterial community (more details in the Result section, Figure 1). The strains were either inoculated alone on seeds or as SynComs of 6, 8, or 12 strains in sterile water. Subsamples of 30 seeds were dipped into bacterial suspensions (OD_600nm_ = 0.01, approximately 10^7^ CFU mL^-1^) for 30 minutes under constant agitation (70 rpm) at 20°C. For the SynComs, each strain was added in equal proportion in the bacterial suspension to reach a final concentration of 10^7^ CFU mL^-1^. Control seeds were inoculated with sterile water only. This concentration was selected based on previous seed inoculation experiments conducted in our team (Chesneau et al. 2022) and multiple reports of this concentration used in synthetic community inoculation on plants (e.g Kwak *et al*. 2018; Pfeilmeier *et al*. 2021). The inoculated or control seeds were then placed individually at the surface of a sterile cotton pad moistened with 4 mL of sterile water in sterile glass tubes. A total of 30 seeds were sown by condition (control, single strains or SynComs, total of 16 conditions) that were incubated for 4 days in a growth chamber (photoperiod: 16h/8h, temperature 25/22°C).

**Figure 1:**
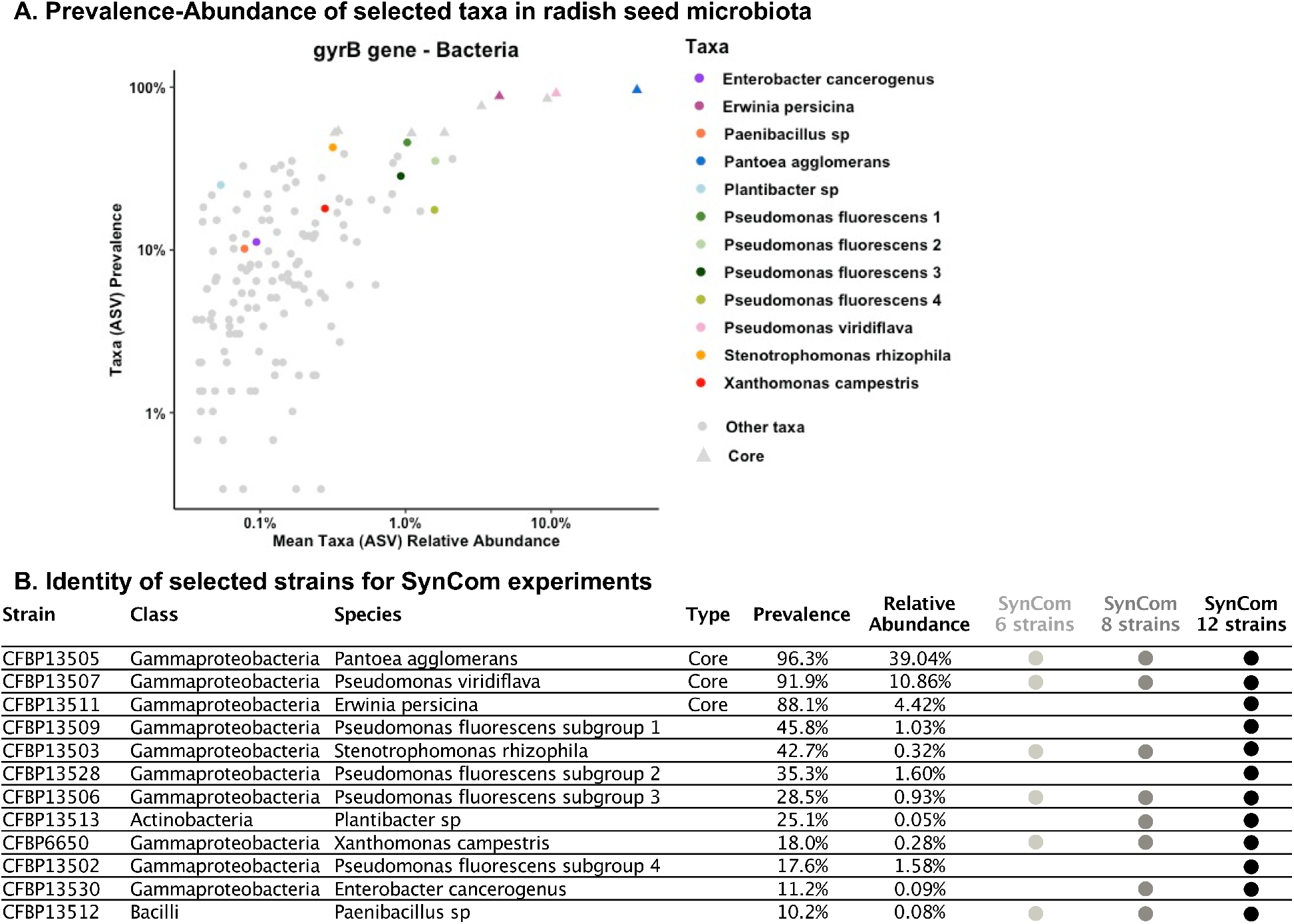
Selection of bacterial strains representative of the radish seed microbiota. A. Prevalence and relative abundance of bacterial ASVs (*gyrB* amplicon sequencing) in radish seed microbiota (*R. sativus* Flamboyant5) from seven independent studies. The colored points represent the taxa selected in this study and the triangle shapes represent the taxa identified as members of the radish core seed microbiota. B. Identity of the strains isolated from radish seeds and used in the SynCom experiments.

The bacterial cell density of the inocula and inoculated seeds (pool of 30 seeds) were assessed by plating on 10% TSA (CFU mL^-1^ or /seed) immediately after inoculating the seeds. In the SynCom experiment, an amplicon sequencing approach was used to measure the relative abundance of the different strains in the inocula, inoculated seeds and seedlings. Thus, 3 repetitions of 500 µL of the inocula and of the macerate of inoculated seeds were stored at -80°C in a 96-well plate before DNA extraction and *gyrB* amplicon library preparation. After 4 days of growth, the phenotypes of the seedlings were determined and then the seedlings were individually processed to assess bacterial cell density by plating and bacterial community composition by amplicon sequencing. The effects on seedling phenotypes were assessed using the protocol established by the International Seed Testing Association (https://www.seedtest.org/en/handbooks-calibration-samples/seedling-evaluation-4th-edition-2018-product-1016.html). Three types of phenotypes could be observed: non-germinated seeds, normal seedling or abnormal seedling. A seedling was considered abnormal if at least one of the cotyledons was necrotic or rotten, if the hypocotyl or epicotyl were deformed, or if the root system was absent, stunted or rotten.

To measure the bacterial cell density and the bacterial communities of seedlings, each entire plant (shoot and root together) without the remaining seed and tegument was crushed in a sterile plastic bag (n=30 seedlings by condition), resuspended in 2 mL of sterile water and homogenized. The seedling suspension was then serial-diluted and plated on 10% TSA to determine the CFU seedling^-1^ and to assess the capacity of strains and SynComs to colonize seedlings. For graphical representations, we calculated the log values of bacterial population density for each sample separately and then calculated the descriptive statistics of each condition to represent them as box plots. The remaining seedling suspensions were stored at -80°C in a 96-well plate before DNA extraction and library preparation for the *gyrB* gene amplicon sequencing.

### Amplicon sequencing of seed and seedling bacterial communities

DNA extraction was performed on the inocula, inoculated seeds and seedlings of the SynCom conditions (control, 6-strain, 8-strain, 12-strain SynComs) with the NucleoSpin® 96 Food kit (Macherey-Nagel, Düren, Germany) following the manufacturer’s instructions.

The first PCR was performed with the primers gyrB_aF64/gyrB_aR553 (Barret *et al*. 2015), which target a portion of *gyrB* gene in bacteria. PCR reactions were performed with a high-fidelity Taq DNA polymerase (AccuPrimeTM Taq DNA Polymerase High Fidelity, Invitrogen, Carlsbad, California, USA) using 5µL of 10X Buffer, 1µL of forward and reverse primers (100µM), 0.2µL of Taq and 5 µl of DNA. PCR cycling conditions constituted an initial denaturation step at 94°C for 3 min, followed by 35 cycles of amplification at 94°C (30 s), 55°C (45 s) and 68°C (90 s), and a final elongation at 68°C for 10 min. Amplicons were purified with magnetic beads (Sera-MagTM, Merck, Kenilworth, New Jersey). The second PCR was conducted to incorporate Illumina adapters and barcodes. The PCR cycling conditions were: denaturation at 94°C (1 min), 12 cycles at 94°C (1 min), 55°C (1 min) and 68°C (1 min), and a final elongation at 68°C for 10 min. Amplicons were purified with magnetic beads and pooled. Concentration of the pool was measured with quantitative PCR (KAPA Library Quantification Kit, Roche, Basel, Switzerland). Amplicon libraries were mixed with 5% PhiX and sequenced with MiSeq reagent kits v2 500 cycles (Illumina, San Diego, California, USA). A blank extraction kit control, a PCR-negative control and PCR-positive control (*Lactococcus piscium*, a fish pathogen that is not plant-associated) were included in each PCR plate. The raw amplicon sequencing data are available on the European Nucleotide Archive (ENA) with the accession number PRJEB58635.

The bioinformatic processing of the amplicons was performed in R. In brief, primer sequences were removed with cutadapt 2.7 (Martin 2011) and trimmed fastq files were processed with DADA2 version 1.10 (Callahan *et al*. 2016). Chimeric sequences were identified and removed with the removeBimeraDenovo function of DADA2. Amplicon Sequence Variant (ASV) taxonomic affiliations were performed with a naive Bayesian classifier (Wang *et al*. 2007) with our in-house *gyrB* database (train_set_gyrB_v4.fa.gz) available upon request. Unassigned sequences at the phylum level and *parE* sequences (a *gyrB* paralog) were filtered. The identification of sequence contaminants was assessed using decontam version 1.8.0 based on our negative controls. After these filtering steps, samples with less than 1000 reads were excluded from the study. The index hopping rate was assessed using our positive controls and the values for this sequencing run were < 0.01% as expected for the Illumina platform.

### Statistics and microbial community analyses

All the scripts and datasets used to conduct the study are available on GitHub (https://github.com/marie-simonin/Radish_SynCom). We assessed the effect of the inoculation and sample type on the various univariate variables (e.g. seedling bacterial cell density, strain relative abundance, ASV richness) using generalized mixed models (*glm* function in lme4 package) and post hoc comparisons were performed using the Tukey method (*warp*.*emm* function in package *emmeans*). The statistical tests performed to compare the number of normal and abnormal seedling phenotypes between control and inoculated seeds (single strain or SynCom) was done using a Fisher exact test for count data.

The analyses on bacterial community structure based on the amplicon sequencing data were done after rarefaction at 14116 reads per sample to conserve sufficient replicates while having a good sampling depth. Diversity and community structure analyses were performed in R 3.6.2 using the *phyloseq* (v1.28.0), *vegan* (v2.5-7) and *microbiome* (v1.7.21) packages (Oksanen *et al*. 2007; McMurdie and Holmes 2013; Lahti, Shetty and Blake 2017). The diversity of the inocula, inoculated seeds and seedlings was characterized using ASV richness. The effects of inoculation and sample types on bacterial community structure were assessed using Bray-Curtis dissimilarity associated with a permutational multivariate analysis of variance (*adonis and pairwise*.*adonis* functions, 999 permutations). Non-Metric Multidimensional Scaling (NMDS) were used to plot the ordinations. The influence of the inoculation of community beta-dispersion was assessed using the *betadisper* function in *vegan* based on Bray-Curtis distance matrix and distance to centroid to each condition. The statistical effects were evaluated using a permutation-based test of multivariate homogeneity of group dispersions followed by a Tukey’s honest significant differences test.

All figures were prepared using the *ggplot2* (v3.3.3) package and the data management was done using the *dplyr* (v1.0.4) and *tidyverse* (v1.3.0) packages in R.

## RESULTS

### 1. Bacterial strain selection based on a meta-analysis of radish seed microbiota

A bacterial culture collection isolated form radish seeds composed of 528 strains was obtained. The strains represented the 4 main phyla (Proteobacteria, Bacteroidetes, Firmicutes, Actinobacteria) found in radish seeds, for a total of 11 assigned bacterial families and 17 genera. For this study, we have selected 12 bacterial strains representatives of the diversity and prevalence-abundance profiles of radish seed microbiota (Figure 1A). These strains were selected without any *a priori* on their potential functions or effects on plant phenotype since seed bacterial communities include a diversity of plant-pathogens, commensals and mutualistic taxa (e.g. Simonin *et al*. 2022). The selection included three (out of nine) extremely abundant and prevalent taxa (i.e core taxa, prevalence >80%, Figure 1A and 1B), but also five sub-dominant (relative abundance < 1.5% and prevalence < 20%) and four rare taxa (prevalence < 20%). Ten strains belonged to Gammaproteobacteria (Enterobacteriaceae, Erwiniaceae, Pseudomonadaceae, Xanthomonadaceae) and the two remaining strains were from the Actinobacteria and Bacilli classes (Figure 1B). In addition to selecting strains that were phylogenetically diverse, we also included infra-species diversity with four strains of the *Pseudomonas fluorescens* subgroup (Hesse et al., 2018) with contrasted prevalence and abundance profiles. Indeed, 380 ASVs affiliated to *Pseudomonas* (25% of the total bacterial richness and 33% of the bacterial relative abundance) were previously detected in the seven radish datasets used in the meta-analysis (Simonin *et al*. 2022). Among this genus, the most diverse species group was *Pseudomonas fluorescens* with 62 ASVs that justified its selection to study interactions between closely related species. The twelve strains selected presented different *gyrB* gene sequences (subspecies level detection) to enable their individual tracking using *gyrB* amplicon sequencing. The *gyrB* sequences of the selected strains and of all bacteria detected in the study (seeds and seedlings) are available in Supplementary Data. The strains were studied in isolation and in SynComs of 6, 8 and 12 strains (Figure 1B) to match the bacterial diversity observed on individual radish seeds (median = 8 ASVs; Figure S1; Chesneau et al. 2022). We performed a nested design for the three diversity levels, with the 6-strain SynCom composed of two high prevalence strains, two intermediate and two low prevalence strains. The 8-strain and 12-strain SynComs were built on this initial community with an increasing number of high-, intermediate- and low-prevalence strains.

### 2. Transmission of single strains and synthetic communities from seed to seedling

#### a. Bacterial colonization of seedlings by all strains and SynComs

Transmission of the twelve individual strains and the three SynComs to radish seedlings was first assessed. When individual strains were seed-inoculated, they were all successfully transmitted to seedlings with a median of 6.8 log CFU seedling^-1^. The seedlings originating from seeds inoculated with SynComs were colonized at similar levels of bacterial cell density (median = 7.4 log CFU seedling^-1^). Single strains and SynComs all reached around 7 log CFU seedling^-1^ that appear to be the bacterial carrying capacity of a radish seedling in our system. The bacterial densities on the control seedlings that originated from surface-sterilized seeds were below detection limit or close to 2 log CFU seedling^-1^, validating that the bacterial populations observed on seedlings in the different conditions originated from our inocula.

#### b. Bacterial community structure of seeds and seedlings inoculated with SynComs

The next step was to assess our ability to manipulate the seed and seedling bacterial community using SynCom inoculation compared to non-inoculated control seeds. Using *gyrB* gene amplicon sequencing, it was validated that the SynCom inoculation led to a significant modification of seed and seedling bacterial community compared to the control condition (pairwise adonis: P<0.007; Figure 3A). On the NMDS ordination, the three sample types (inoculum, seed and seedling) clustered closely by SynCom indicating a high degree of similarity, whereas the control seeds and seedlings were very dispersed. This observation was further confirmed by a beta-dispersion analysis (distance to centroid) indicating that the SynCom inoculation strongly reduced the variability in seed and seedling community structure compared to the control condition (Figure 3B, 3C). Furthermore, it was confirmed that the three SynComs were significantly distinct in the three types of samples (inocula, seeds, seedlings, Figure 3D), with an expected structuration showing the 6 and 12-strain SynComs as the most different and the 8-strain SynCom with an intermediate composition.

**Figure 2:**
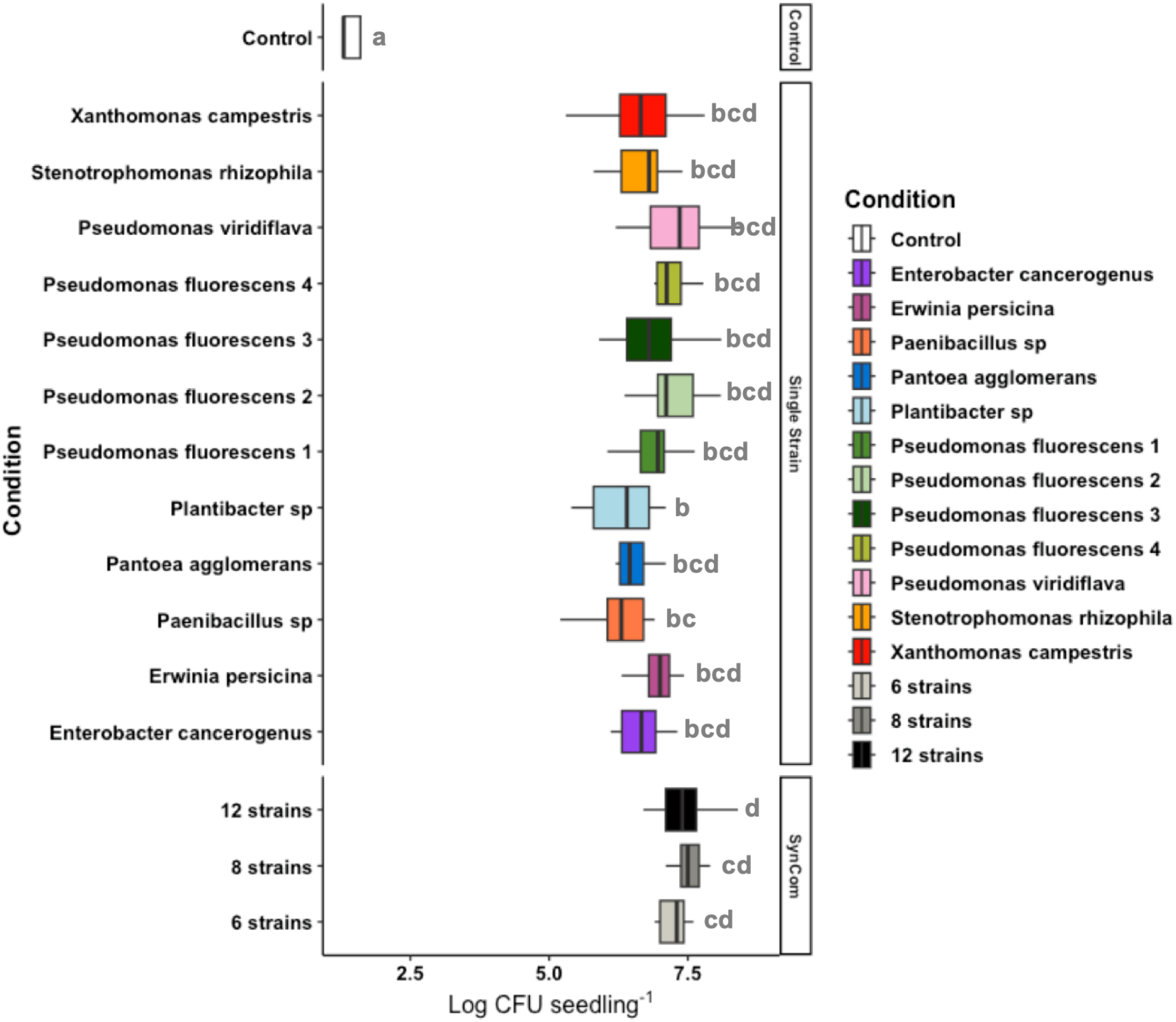
High seedling bacterial colonization (bacterial population densities > Log 7 CFU seedling^-1^) in the different inoculation treatments (n=30 seedlings per condition). CFU: Colony Forming Unit. The different letters represent the results of a post-hoc Tukey HSD test.

**Figure 3:**
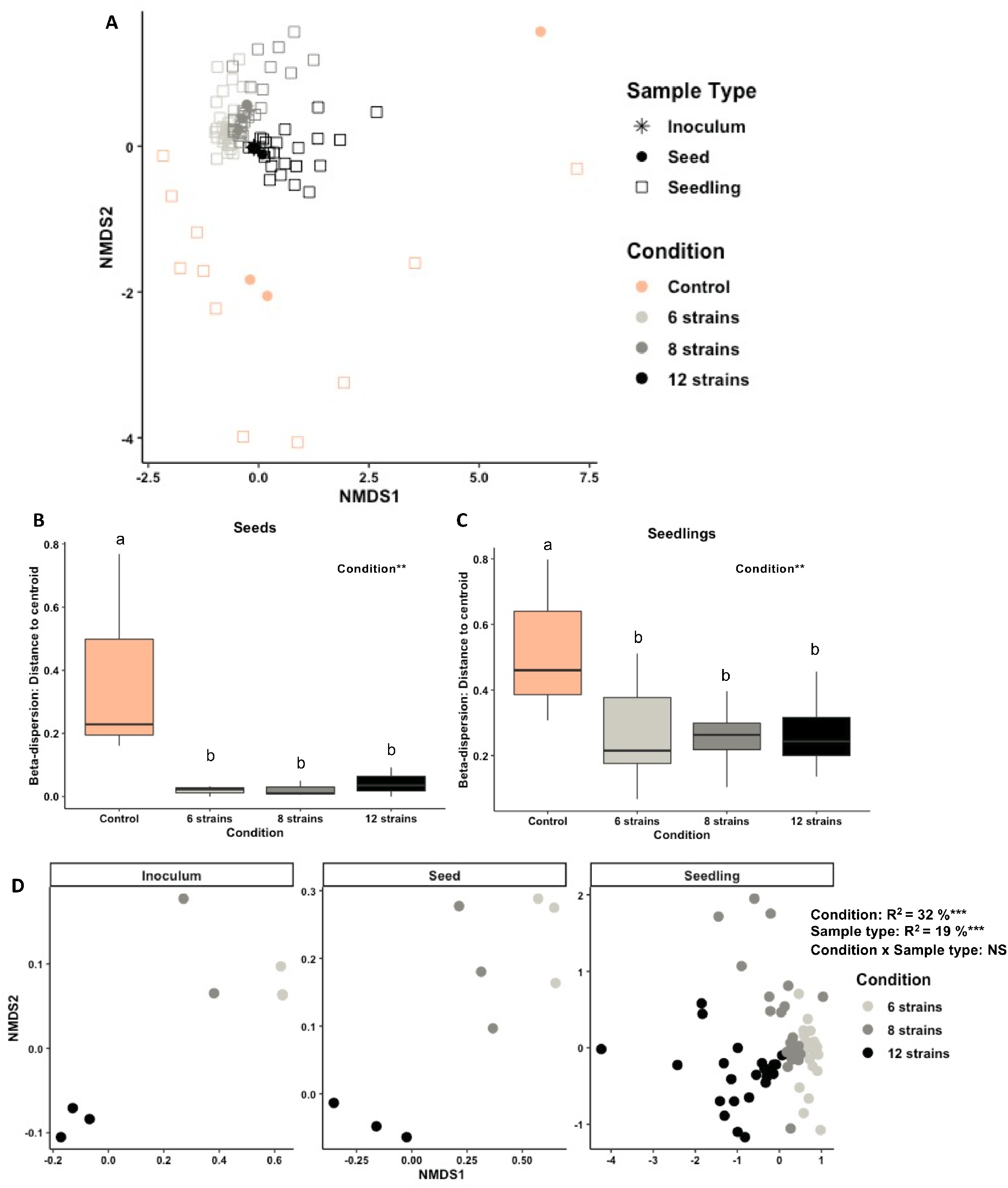
Impact of SynCom inoculation on seed and seedling bacterial community structure. A) Bacterial community structure for the three sample types and inoculation treatments visualized through an NMDS ordination based on Bray-Curtis distances (stress = 0.155). Beta-dispersion assessed using distance to centroid of the B) seed and C) seedling community structure. D) Bacterial community structure of the three SynComs in the inocula, seeds and seedlings visualized through an NMDS ordination based on Bray-Curtis distances (stress = 0.165). The asterisks represent the significance of the Permanova test: \*\*\**P*<0.001, NS=Non-Significant. The different letters represent the results of a post-hoc Tukey HSD test; two conditions with no letters in common are statistically different.

#### c. Effect of SynCom inoculation on seed and seedling richness

Next, the effect of SynCom inoculation on the bacterial richness of seeds and seedlings was assessed. It was confirmed that SynCom inoculation enabled the reconstruction of a diversity gradient on both seeds and seedlings (Figure 4A). The bacterial richness measured on seeds and seedlings corresponded to the expected taxa richness present in the inocula (6, 8 or 12 ASVs). The control seeds were surface-disinfected but they still harbored a low bacterial diversity likely to be endophytic bacteria, including ASVs of strains included in the SynComs because the strains selected have been isolated from the same radish genotype. The microbiota comparison of native and surface-disinfected seeds suggests that seed endophytes still present after disinfection are dominated by *Pseudomonas* species and *Pantoea agglomerans* (Figure S2). Altogether, these results show the efficacy of the SynCom inoculation on seeds to drive the bacterial diversity of both seeds and seedlings.

**Figure 4:**
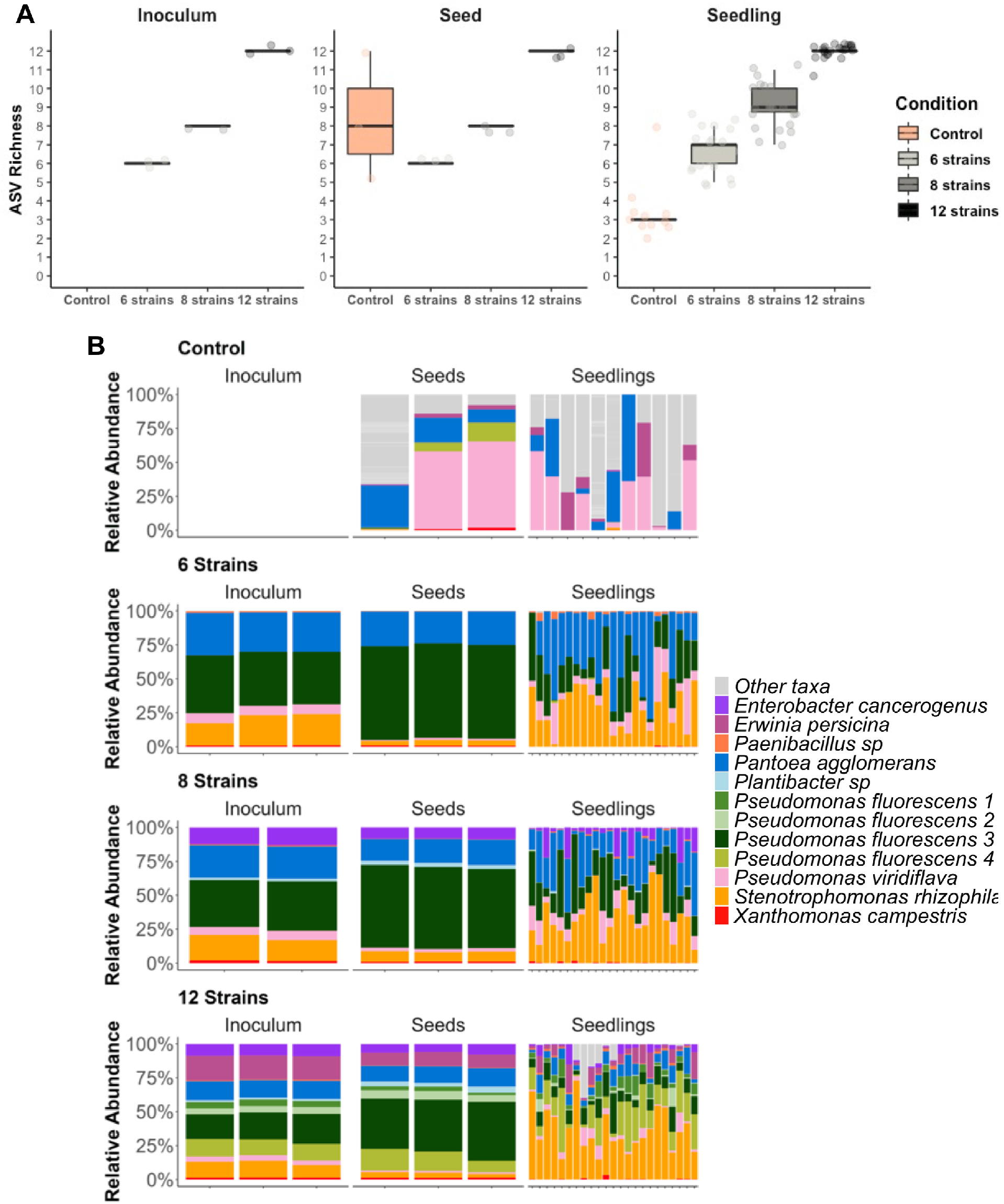
Shifts in bacterial diversity and composition of seeds and seedlings inoculated with SynComs. A) Bacterial taxa richness (ASV richness) of the inocula, inoculated seeds and seedlings in the different conditions. B) Taxonomic profile of the inoculum, seeds and seedlings in the four different conditions (control, 6 strains, 8 strains and 12 strains). Each stacked bar represents a sample. The composition of the “other taxa” in control seeds and seedlings are presented in Figure S2 and S3, respectively.

#### d. Tracking of the transmission of the SynCom strains between the inoculum, seeds and seedlings

Based on the unique *gyrB* ASV of the 12 strains, we were able to track their transmission patterns from the inoculum to the seedling. First, we characterized the natural abundance of the strain ASVs in the control seeds and seedlings. The 12 ASVs were naturally found in control seeds and seedlings, especially the ASVs associated with the *Pseudomonas viridiflava, Pantoea agglomerans, Pseudomonas fluorescens* 4 and *Erwinia persicina* strains (Figure 4B). A complete taxonomic description of the control seedlings (including the “other taxa”) is available in Figure S3. It shows that control seedling bacterial community is highly variable between individuals and is generally dominated by 2 to 9 ASVs. As already seen in Figure 3A, the SynCom inoculations led to completely different seed and seedling bacterial community compositions than in the control condition (Figure 4B). Interestingly, we observed some differences in the abundance of strains between the inoculum and the seed community profile (Figure 5A), indicating differences between strains for their capacity to colonize the seed during inoculation (Figure 5B). For instance, the strain *Pseudomonas fluorescens* 3 had a higher relative abundance on seeds (40-70%) compared to the inocula (20-40%), while the strains *Pseudomonas viridiflava* and *Stenotrophomonas rhizophila* had the opposite pattern (Figure 5B).

**Figure 5:**
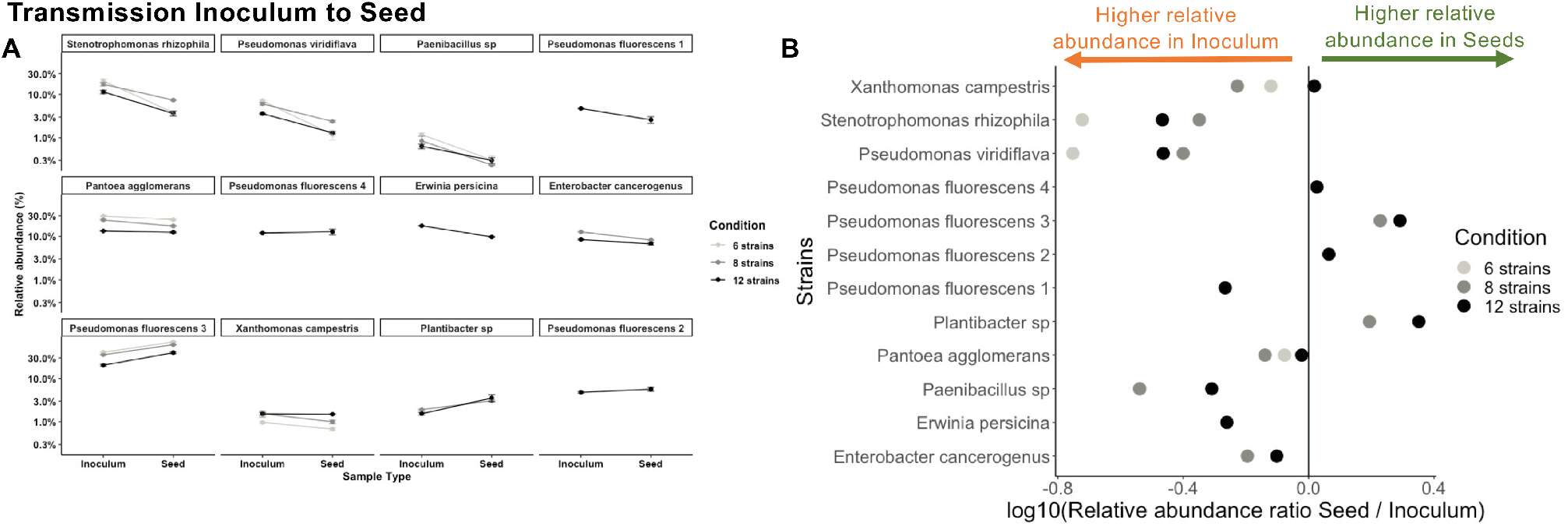
Contrasted capacities of strains in seed colonization A. Relative abundance patterns of each strain during the transmission from inoculum to seed (A) in synthetic bacterial communities. B. Ability of each strain to colonize seeds compared to their initial relative abundance in the inocula: ratio of relative abundance seed / inoculum.

During the phenological transition from seed to seedling, we observed an important restructuration of the bacterial community (Figure 4B). All the strains were able to transmit from seeds to seedlings but with variable success depending on the SynCom richness and the seedling individual (Figure 4B, 6). Across the three SynComs, *Stenotrophomonas rhizophila* was the most successful seedling colonizer (mean relative abundance = 30 to 36%), followed by *Pantoea agglomerans* but only in the 6- and 8-strain SynComs (mean relative abundance = 34 and 25%). In contrast, the strain *Pseudomonas fluorescens* 3 that was dominant on seeds strongly decreased in abundance on seedlings (11 to 24%).

The fate of each strain was monitored during the seed to seedling transmission across the three SynComs and we grouped the strains in three categories: increasers, stable, decreasers (Figure 6A). Four ‘increaser’ strains were identified and three of them presented this positive transmission pattern across the three SynComs (*Stenotrophomonas rhizophila, Pseudomonas viridiflava, Paenibacillus sp*). In contrast, four ‘decreaser’ strains were identified with three of them with a consistent negative transmission pattern (*Pseudomonas fluorescens* 3, *Xanthomonas campestris, Plantibacter sp*.). Finally, four strains presented a stable transmission or a variable profile depending on the inoculated SynCom. For instance, *Pantoea agglomerans* and *Enterobacter cancerogenus* had a stable transmission in the 6- or 8-strain SynComs, but decreased in the 12-strain SynCom. Interestingly, the 4 strains of *Pseudomonas fluorescens* tested presented contrasted transmission patterns or abundance profiles and were found in the three categories (increaser, stable or decreaser). Another interesting result was that the four strains identified as “increasers” were not the best seed colonizers after inoculation (Figure 5A, 5B). Inversely, some of the best seed colonizers (*Pseudomonas fluorescens* 3) were identified as “decreasers” on seedlings, indicating contrasted strain fitness depending on the plant phenological stage.

**Figure 6:**
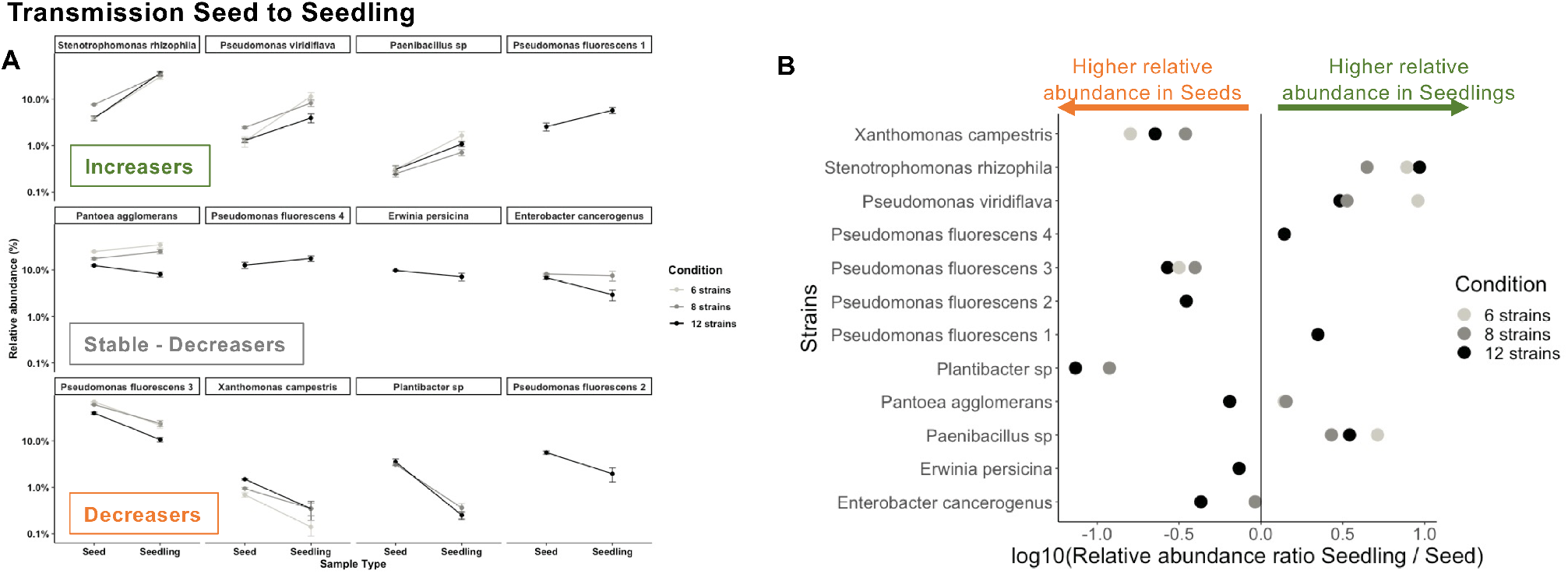
Contrasted capacities of strains in seedling colonization A. Relative abundance patterns of each strain during the transmission from seed to seedling. B. Ability of each strain to colonize seedlings compared to their initial relative abundance on seeds: ratio of relative abundance seedling / seed.

We also analyzed the influence of the SynCom composition on the ability of each strain to colonize seedlings compared to their initial relative abundance on seeds (i.e ratio of relative abundance seedling / seed, Figure 6B). We observed that the SynCom richness strongly influenced the abundance patterns of the strains. Most strains had a lower seedling colonization in the 12-strain SynCom (e.g. *Pantoea agglomerans, Enterobacter cancerogenus*), with the exception of *Stenotrophomonas rhizophila* (Figure 6B). Some strains had a higher seedling colonization in the 6-strain SynCom than the 8-strain SynCom (*Pseudomonas viridiflava, Paenibacillus sp, Stenotrophomonas rhizophila*). Altogether, these results indicate strong fitness differences between strains in the seed and seedling habitat, that is highly dependent on the surrounding community.

### 3. Effect of single bacterial strains or synthetic communities on seedling phenotype

#### a. Identification of strains and SynComs increasing the proportion of abnormal seedlings and non-germinated seeds

The impact of the inoculation of single strains or SynComs on seedling phenotypes was assessed based on the proportion of non-germinated seeds, normal and abnormal seedlings observed (Figure 7). In the control condition, we observed a low proportion of non-germinated (7%) and abnormal seedlings (7%). Several individual strains and the three SynComs caused detrimental effects on seedling phenotypes. In particular, *Pseudomonas viridiflava, Paenibacillus sp* and two of the SynComs (6 and 8 strains) had significantly larger proportions of abnormal seedlings (60 to 73%). The 12-strain SynCom also led to an increased proportion of abnormal seedlings (34%), but to a lesser extent than the other two SynComs. The conditions that presented the highest proportion of normal seedlings were *Stenotrophomonas rhizophila* (90%), *Pseudomonas fluorescens* 3 (90%) and *Plantibacter sp* (87%).

**Figure 7:**
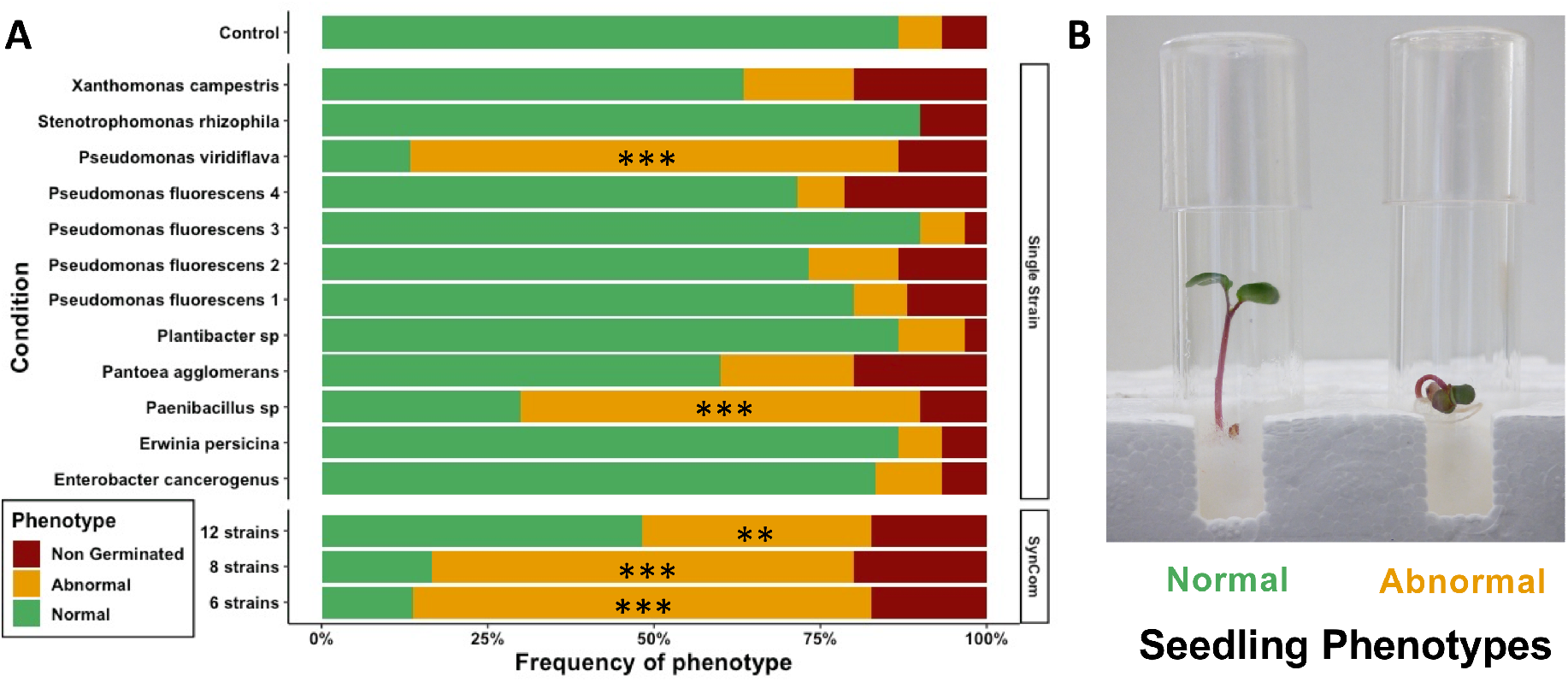
A) Effect of the inoculation of single bacterial strains or synthetic bacterial community on germination and seedling phenotypes. The asterisks represent the significance of the Fisher exact test that compares the proportions of normal and abnormal seedling phenotypes between control and inoculated seeds: \*\*\**P*<0.001, **P<0.01. B) Image of the typical phenotypes observed in the experiment (Credit: Guillaume Chesneau).

#### b. Modifications in the composition of seedling microbiota between normal and abnormal seedlings

Next, we analyzed if the seedling bacterial community structure was different depending on the seedling phenotype (Figure 8A). The seedling phenotype was a significant driver of seedling microbiota (R^2^=3.7%), especially in the 12-strain SynCom (interaction Condition x Phenotype, *P*=0.006), for which the number of normal and abnormal seedlings was more balanced (14 normal vs 10 abnormal). In the 12-strain SynCom, we analyzed if some strains had a significantly higher or lower relative abundance in abnormal seedlings (Figure 8B). The strains *Enterobacter cancerogenus* and *Erwinia persicina* were more abundant in abnormal seedlings, while *Pseudomonas viridiflava* and *Xanthomonas campestris* were less abundant.

**Figure 8:**
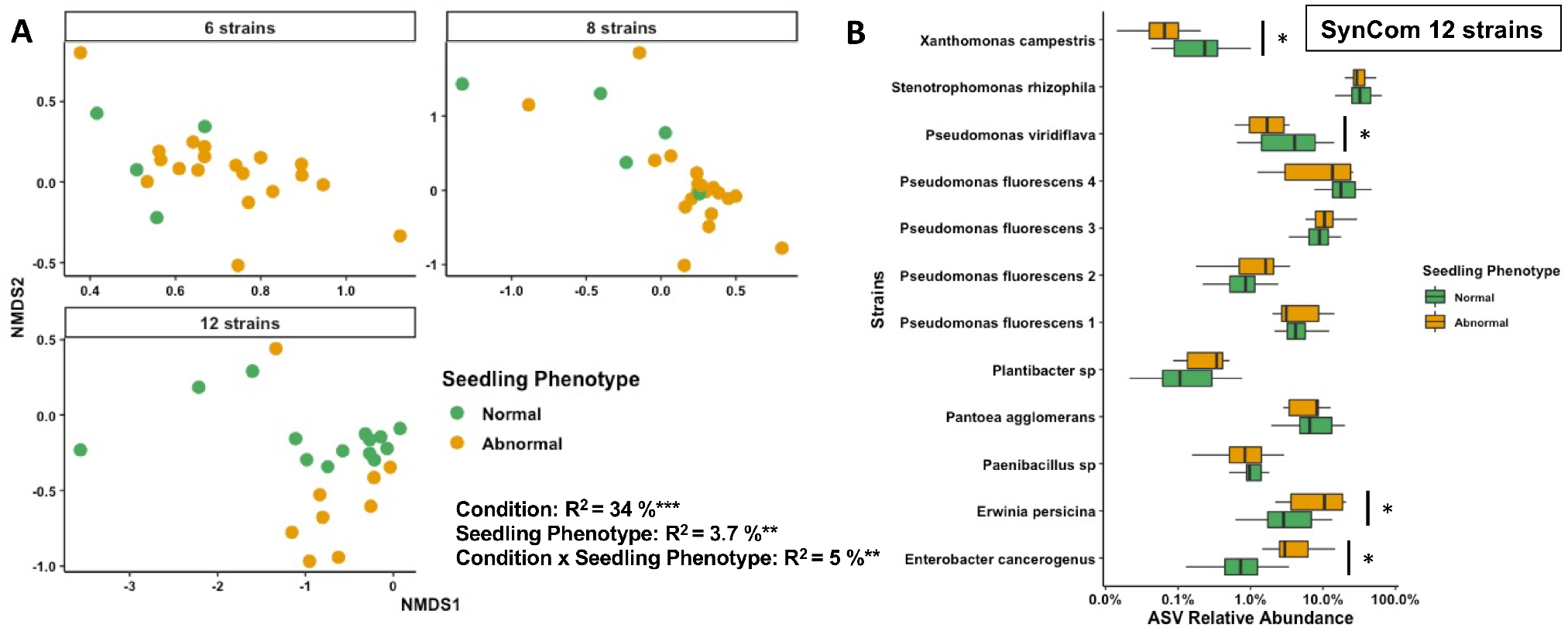
Influence of seedling phenotype (normal or abnormal) on A) bacterial community structure (stress=0.17) and B) on the strain relative abundance on the 12-strain SynCom. The asterisks represent the significance of the Permanova test or post-hoc Tukey test: \**P*<0.05; \*\**P*<0.01; \*\*\**P*<0.001.

## DISCUSSION

### Community- and strain-specific patterns of seedling transmission of seed-borne bacteria

This study demonstrates that a precise manipulation of the seed and seedling bacterial community is possible through SynCom inoculations. The main achievements of this experiment rely on the successful reconstruction of a richness gradient (6, 8 and 12 strains) on both seeds and seedlings and on the strong reduction in the natural variability of microbiota structure (beta-dispersion). SynCom inoculation leads to more homogeneous microbial community compositions between replicates compared to native communities described in Figure S2 or in previous reports from the literature (Moroenyane *et al*. 2021; Walsh *et al*. 2021). This homogenization of community structure is due to the high abundance of inoculated strains (10^7^ CFU seedling^-1^), which makes the members of the native endophytic community (10^2^ CFU seedling^-1^) undetectable. This similarity of seed microbiota between seed replicates is a prerequisite to study the transmission of microorganisms from seed to seedling and establish causal relationship between seed microbiota and plant microbiota or phenotype.

The inoculation of the three different SynComs on seeds led to the assembly of significantly distinct seedling bacterial communities. However, the relative abundance of the strains changed drastically from the inocula to seeds and then to seedlings. Some strains like *Pseudomonas fluorescens* 3 were excellent seed colonizers, while others had a reduced abundance compared to the inocula (e.g *S. rhizophila*). These differences in seed colonization can be due to contrasted adhesion capacity of strains related to the secretion of large adhesins (i.e LapA, LapF) and the presence of flagella (DeFlaun *et al*. 1994; Yousef-Coronado, Travieso and Espinosa-Urgel 2008; Duque *et al*. 2013).

Interestingly, the best seed colonizers were not the best seedling colonizers. In contrast, all the strains that declined during the inocula-seed transition phase increased in the seed-seedling transition (i.e “increasers”). In particular, *S. rhizophila* became dominant on seedlings of the three SynComs. These results indicate that the bacterial traits required to colonize seeds and seedlings are different. Previous studies indeed indicate that seed colonization is associated with high tolerance to desiccation and chemotaxis-induced motility (Truyens *et al*. 2015), while successful seedling colonization is linked to copiotrophic lifestyles with fast growth in response to increased nutrient availability (Barret *et al*. 2015, Torres-Cortes *et al*. 2018). Moreover, these findings suggest that being dominant on seeds does not provide an advantage to become dominant on seedlings. These findings are in line with Rochefort *et al*. (2021) and Chesneau *et al*. (2022) demonstrating a low transmission success of abundant seed taxa to seedlings in natural microbial communities.

Additionally, we observed that SynCom richness influenced the seedling transmission of the strains, indicating an important role of biotic interactions in seedling microbiota assembly. For instance, *Pantoea agglomerans* had a positive seedling colonization success in the 6- and 8-strain SynComs and a negative one in the 12-strain SynCom. These SynCom-specific patterns of each strain are likely driven by exploitative and interference competition between individuals that necessitate subsequent drop-out SynCom experiments to identify the specific mechanisms expressed *in planta* (Hibbing *et al*. 2010; Granato, Meiller-Legrand and Foster 2019).

Our experiments included four different strains of *Pseudomonas fluorescens* to compare the intra-specific patterns of seedling transmission. Both seed and seedling colonization patterns were highly strain-specific. We observed all possible outcomes depending on the *Pseudomonas fluorescens* strain considered (increasers, stable or decreasers). This observation is in line with another study on diverse *Pseudomonas fluorescens* strains that greatly varied in their ability to colonize *Arabidopsis thaliana* alone or in competition with another *Pseudomonas sp*. strain (Wang *et al*. 2022). This result suggests important differences between strains in traits involved in habitat colonization and competitive interactions, that prevent any generalization of these observations at the species level.

Altogether, these results show that SynCom inoculation can effectively manipulate seed and seedling microbiota diversity and highlight strong fitness differences between native seed-borne taxa in the colonization of these habitats.

### Contrasting effects of seed-borne bacteria on seedling phenotype in isolation and in a community context

This study demonstrates the value to study plant phenotype modulation by bacteria in a community setting using SynComs. The proportion of abnormal seedling phenotypes decreased at the highest SynCom richness level (12 strains). This observation could be explained by a “dilution” of the negative effects of the detrimental strains as the diversity increased and their relative abundances decreased. Still surprisingly, the strains enriched in abnormal seedlings in the 12-strain SynCom had not been identified as detrimental in isolation (*Enterobacter cancerogenus* and *Erwinia persicina*). Furthermore, we found that the detrimental *Pseudomonas viridiflava* and the seed-borne pathogen *Xanthomonas campestris* pv. *campestris* were even reduced in abundance in abnormal seedlings in this condition. We can propose different hypotheses to explain these results. One is that the identity of the strains responsible for the abnormal phenotypes are different when inoculated in a community (*Enterobacter cancerogenus* and *Erwinia persicina)* or in isolation (*Pseudomonas viridiflava* and *Paenibacillus sp*.). Another hypothesis is that *Enterobacter cancerogenus* and *Erwinia persicina* are opportunistic taxa that are enriched in abnormal seedlings (e.g necrotrophy or “cry for help” of the plant) but are not responsible for the altered phenotype. Previous reports on diseased plants caused by a known pathogen support these hypotheses as the increase in abundance of beneficial protective strains or opportunistic copiotrophic taxa is frequently observed (Masson *et al*. 2020, Pfeilmeier *et al*. 2021, Arnault *et al*. 2022).

In isolation *Pseudomonas viridiflava* and *Paenibacillus sp* strongly increased the proportion of abnormal seedlings. *Pseudomonas viridiflava* is a very common plant-associated bacterium with various life-styles (endophytes, epiphytes, saprotrophs, pathogens) reported to be extremely prevalent in seeds (Simonin *et al*. 2022) and in the phyllosphere of a large diversity of hosts (Karasov *et al*. 2018, Lipps and Samac 2022, Lundberg *et al*. 2022). *Pseudomonas viridiflava* is frequently seed-transmitted and can cause germination arrest or symptoms on seedlings, including in radish (Shakya and Vinther 1989; Samad *et al*. 2017). *Paenibacillus* strains are mainly recognized for their biocontrol capacities against various bacterial or fungal pathogens (Ryu *et al*. 2006; Ling *et al*. 2011), but one report indicates root damages caused by this bacteria under gnotobiotic conditions like in our study (Rybakova *et al*. 2016). Another important observation was that the three bacterial strains identified as seed core taxa (*Pseudomonas viridiflava, Pantoea agglomerans, Erwinia persicina)* were associated with detrimental effects on seedling phenotypes either in isolation or in SynComs. These results are in line with other studies showing that the plant core microbiota includes pathogens or conditional pathogens that have a pathogenic potential but are generally kept in check by other members of the microbiota and the host (Pfeilmeier *et al*. 2021, Simonin *et al*. 2020).

Additionally, we show that the inoculation of multiple strains with potential pathogenic effects (i.e pathobiome) do not have additive effects that could have led to an absence of germination or 100% of abnormal seedlings. The reduction of detrimental effects in the 12-strain SynCom experimentally confirms the importance of bacterial interactions in plant disease development in a pathobiome context (Bass *et al*. 2019). This observation is supported by other SynCom experiments on the phyllosphere demonstrating the role of single or consortia of commensal microorganisms (e.g *Pseudomonas*) in disease protection (Vogel *et al*. 2021, Shalev *et al*. 2022). Increasing microbial interactions with higher SynCom diversity is a promising leverage for reducing the impact of deleterious bacterial strains.

Altogether, these results indicate that this synthetic ecology approach is valuable to better understand the assembly of plant microbiota in early life stages and the context dependency of disease expression in a community setting. This approach permitted identifying several seed core bacteria with detrimental effects on germination and seedlings but also to characterize the transmission success of diverse bacterial strains (i.e increasers, stable, decreasers) to seedlings in three community contexts.

## ACKNOWLEDGEMENTS

The authors are grateful to Guillaume Chesneau, Gloria Torres-Cortes and Samir Rezki that produced several radish seed microbiota datasets used in the meta-analysis performed. We thank Emmanuelle Laurent and Vincent Odeau (FNAMS) for field crop management and the production of the initial seed samples used in this study. We thank Muriel Bahut (ANAN platform, SFR QuaSav) for amplicon sequencing. We also thank the International Centre of Microbial Resource (CIRM) - French Collection for Plant-associated Bacteria (CFBP – INRAE, https://doi.org/10.15454/E8XX-4Z18) for the conservation of the bacterial strains of the study. The authors declare that they have no conflicts of interest.

## FUNDING

This work was supported by the French National Research Agency [ANR-17-CE20-0009-01].

## DATA AVAILABILITY

The raw amplicon sequencing data are available on the European Nucleotide Archive (ENA) with the accession number PRJEB58635.

All the scripts (bioinformatics, analyses, figures) and datasets used to conduct the study are available on GitHub (https://github.com/marie-simonin/Radish_SynCom).

## SUPPLEMENTARY INFORMATION

**Figure S1:**
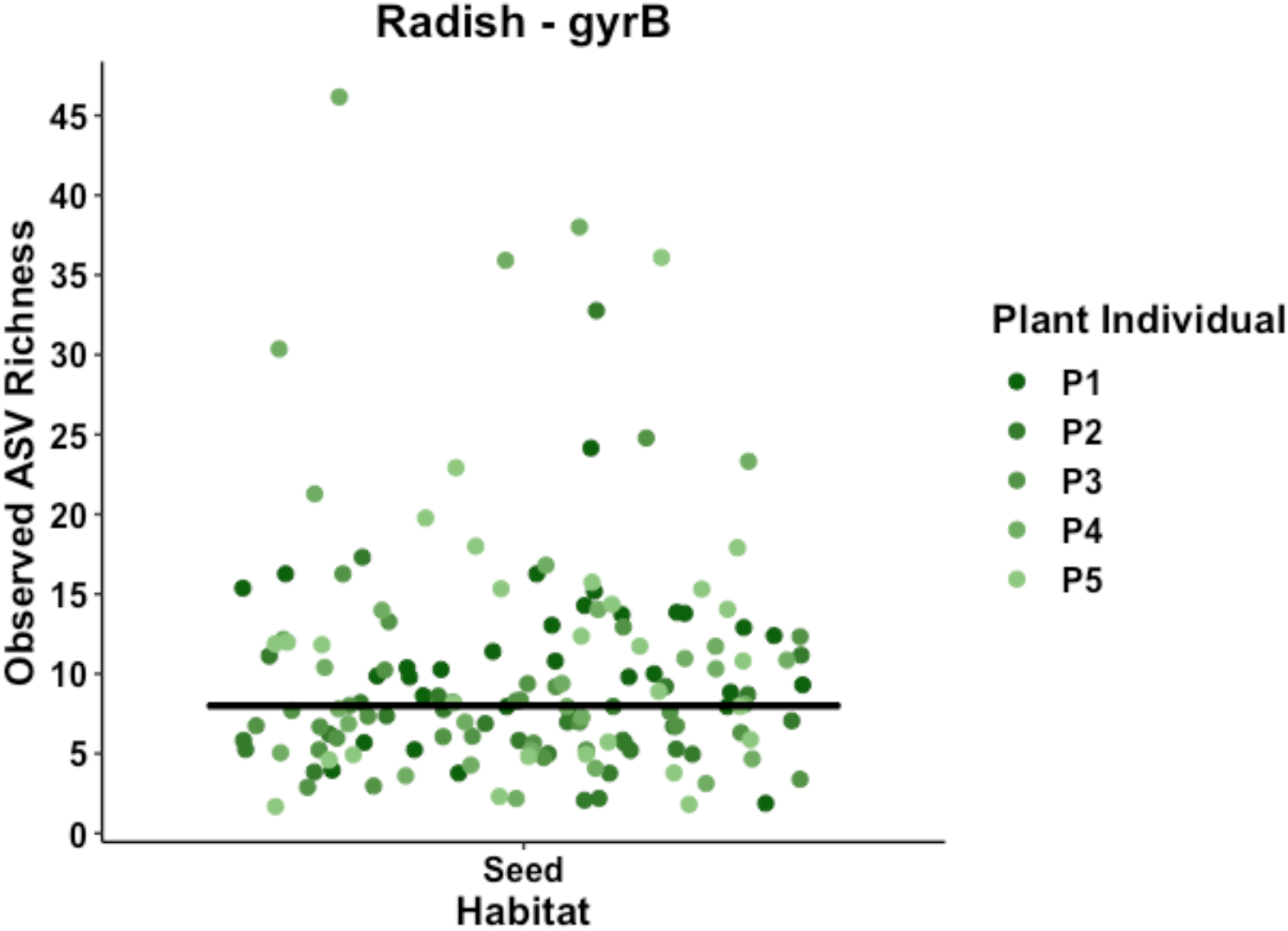
Observed ASV richness of individual mature seeds of radish (Flamboyant5). The data were obtained from Chesneau et al. (2022) based on 149 seeds analyzed. The horizontal line represents the median value at 8 ASVs per seed.

**Figure S2:**
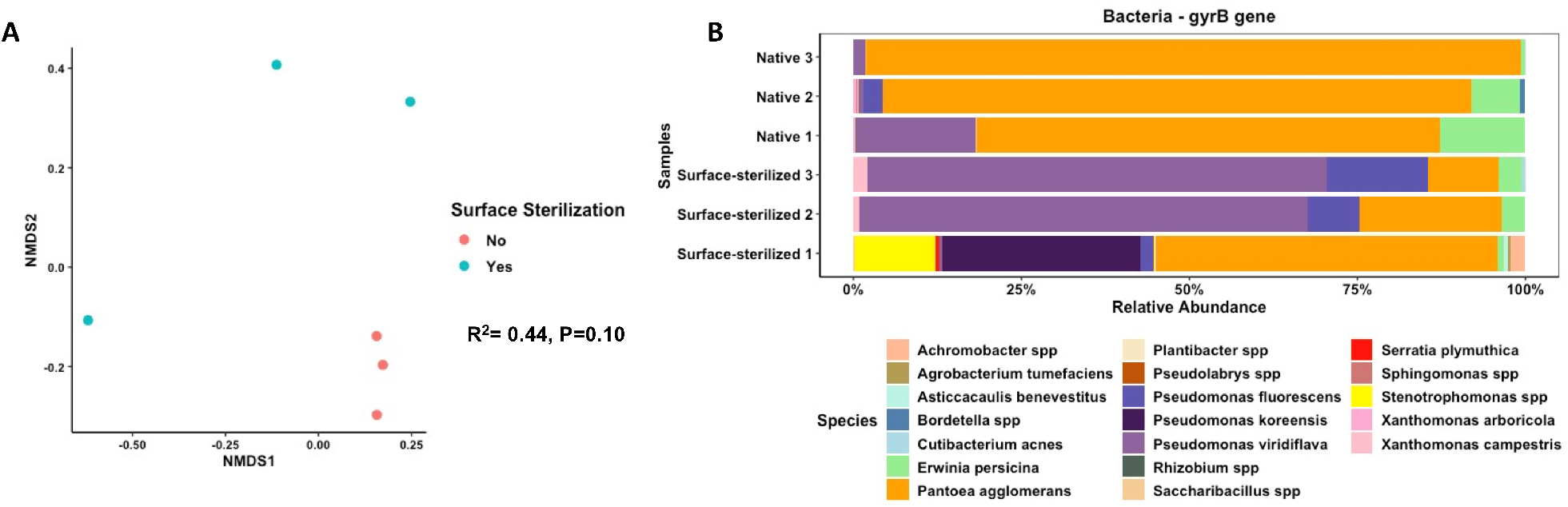
Characterization of the bacterial community of native and surface-sterilized seeds. A. Ordination based on Bray-Curtis distances, B. Taxonomic composition at the species level of each seed sample analyzed.

**Figure S3:**
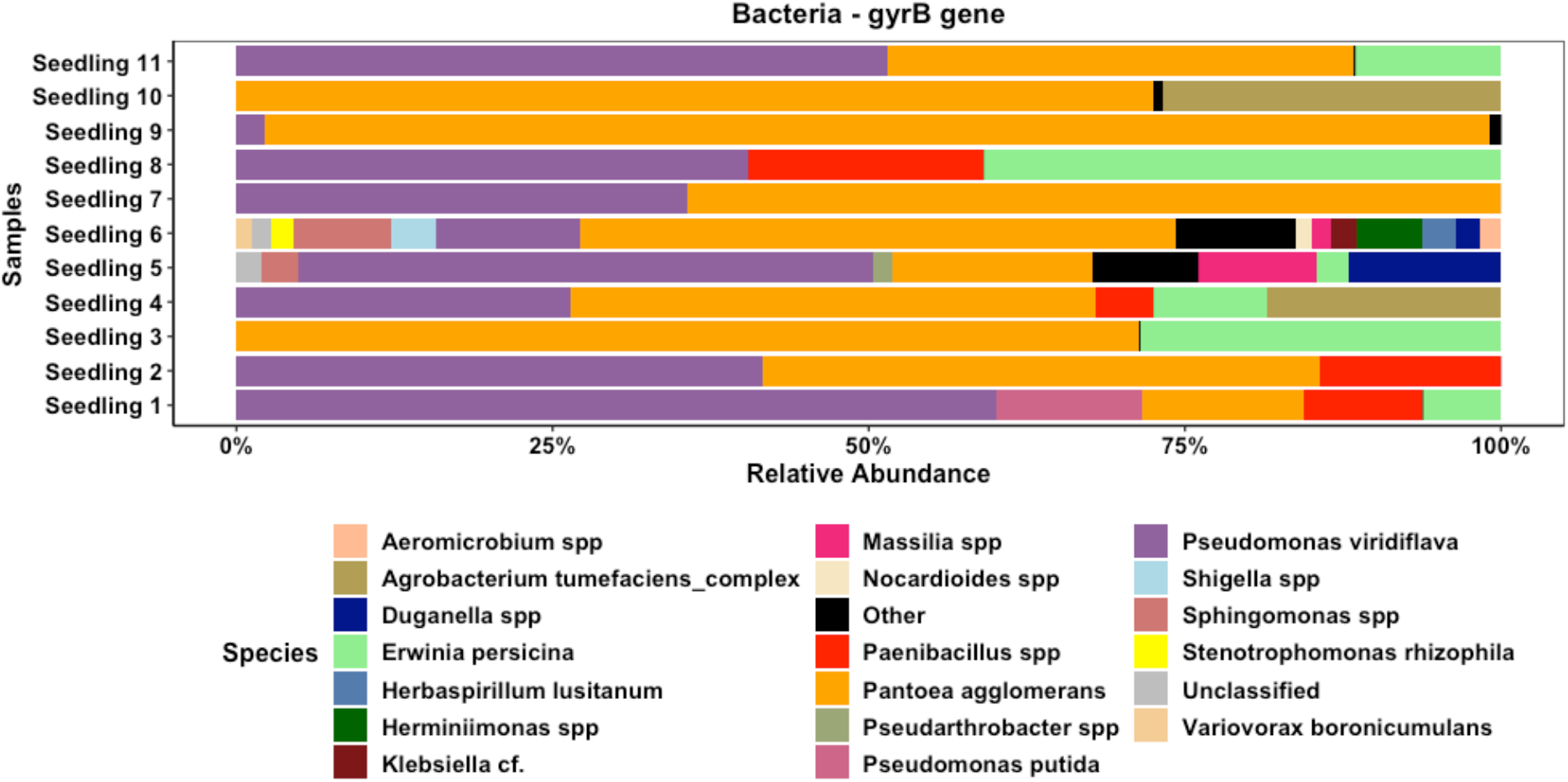
Bacterial taxonomic profile of the control seedlings at the species level. “Other” species represents taxa with a relative abundance inferior to 1%.

## Notes

### Competing Interest Statement

The authors have declared no competing interest.

### Summary of Updates

Second revision of paper following the review of the article by PCI Microbiology recommenders

